# Automated analysis of feeding dynamics from electromyographic recordings in a blood-sucking insect

**DOI:** 10.64898/2026.05.29.728826

**Authors:** Héctor Salas Morales, Isabel Ortega-Insaurralde, Marcelo G. Armentano, Ariel Monteserin, Pablo E. Schilman, Romina B. Barrozo

## Abstract

Feeding behavior in blood-sucking insects relies on gustatory evaluation to decide on sustained ingestion, yet quantifying this process from electromyogram (EMG) recordings is labor-intensive. Here we developed MyoRec, an automated computational framework employing machine learning to analyse EMG signals from the triatomine bug *Rhodnius prolixus*. Using recordings under appetitive and aversive conditions, a convolutional neural network detected ingestion events with 97.7% accuracy. Automated analysis revealed distinct feeding dynamics, with prolonged ingestion and higher pumping frequency under appetitive stimuli, compared to rapid feeding cessation under aversive stimuli. MyoRec substantially reduces analysis time while maintaining accuracy, providing a scalable tool to investigate how gustatory cues modulate feeding decisions in hematophagous insects.

## 1. Introduction

Animals rely on taste to assess the quality of potential food sources and avoid ingesting harmful substances that may cause malaise, illness, or death. The gustatory system is specialized to detect appetitive compounds, such as sugars, amino acids, low concentration of salts, and nucleotides, as well as aversive compounds, including bitter and acidic substances and high concentration of salts [1]. In hematophagous insects, this gustatory evaluation of the ingested fluid is specially critical for decisions to initiate or continue feeding [2,3].

Triatomines are hematophagous insects of medical importance because they serve as vectors of *Trypanosoma cruzi*, the causative agent of Chagas disease [4]. Among them, *Rhodnius prolixus* is one of the most extensively studied species, both due to its health relevance and its long-standing use as a model organism in fields such as biochemistry, physiology, molecular biology, ecology, toxicology, metabolism, among others [5–8].

The feeding behaviour of triatomines consists of two main stages: sampling and ingestion [9,10]. During the sampling phase, the insect ingests a small portion of blood to evaluate its suitability; if the blood contains the appropriate stimulatory compound, such as phagostimulants, the insect proceeds to full ingestion [2,10] This initial sampling phase is a critical behavioral decision point that determines whether feeding will continue or be interrupted.

Understanding feeding behavior is essential for identifying the molecules that promote or inhibit ingestion. In insects, compounds such as nucleotides and other nutrients act as phagostimulants, while deterrent substances suppress ingestion [3,6,11]. Quantifying the effects of these chemical cues on feeding dynamics requires reliable methods for detecting ingestion events and extracting behavioral parameters.

Electromyography (EMG) is a technique that records the electrical activity of the sucking muscles in the insect’s head during feeding [10,12,13]. In this method, insects feed from an artificial feeder or on a live host equipped with electrodes, and the resulting recordings, electromyograms (EMGs), capture waveforms over time that correspond to different phases and behaviors [14].

Like many other insects, triatomines generate suction for feeding through the coordinated activity of two muscle groups in the anterior part of the head: the cibarial pump and the pharyngeal pump. These muscles contract and relax in antiphase to produce the pumping action, and this activity can be readily recorded and quantified using EMG [14].

Extracting behavioral information from EMG recordings requires characterization of the wave patterns associated with specific feeding behaviors. In triatomines, distinct waveforms correspond to the sampling phase, onset of the ingestion, and end of feeding. This characterization allows calculation of parameters such as the number of pumping events, ingestion rate, contact time with the food source, and pumping frequency [10,14]. However, manual identification of these waveforms and calculation of behavioral parameters is time-consuming, labor-intensive, and prone to human error, particularly with large datasets.

Understanding how chemical stimuli influence the transition between sampling and sustained ingestion requires quantitative tools that can detect ingestion events and measure feeding dynamics with high temporal resolution.

The aim of this study was to develop a quantitative framework to characterize the feeding behavior of *R. prolixus* and to determine how chemical cues influence ingestion dynamics. To achieve this, we developed MyoRec, a computational platform that combines machine learning algorithms and automated signal analysis to decode feeding behavior from electromyogram recordings, enabling quantitative investigations of feeding decisions in blood-feeding insects. The software comprises three main modules: (1) Ingestion Event Identifier (IEI), which automatically detects ingestion events; (2) Peak Detector (PD), which extracts EMG signals and calculates feeding behavior variables with optional human supervision; and (3) Data Calculator (DC), which automates the calculation of behavioral parameters using fixed settings, enabling rapid batch analysis of multiple recordings with minimal human intervention.

## 2. Materials and Methods

### 2.1 Insects

Fifth-instar larvae of *Rhodnius prolixus* were used in all experiments. Insects were reared at 28 °C, ambient relative humidity (30 - 60 %) and a 12 h: 12 h L:D light:dark cycle. Feeding experiments were carried out with unfed insects 15 ± 2 days post-ecdysis. Recordings were conducted during the first 0–6 h of the scotophase, when triatomines show peak host-seeking and feeding motivation [15,16]. All animals were handled in accordance with the biosafety guidelines of the Facultad de Ciencias Exactas y Naturales, Universidad de Buenos Aires, Buenos Aires, Argentina.

### 2.2 Electromyographic recording setup

Electromyographic (EMG) recordings were obtained using an artificial feeding system adapted from Pontes et al. [10]. The setup consists of a feeding container (FC) positioned above a shelter container (SC). The FC is a glass chamber (1 cm diameter, 2 cm height) sealed with a latex membrane through which the insect inserts its proboscis to access the feeding solution (Fig. 1). The SC beneath the feeder is covered with mesh to prevent escape. Temperature is maintained at 34 ± 2 °C using a thermostatic bath with water recirculation through silicone tubing connected to the FC. A recording electrode is connected to a metal mesh inside the SC; the reference electrode is immersed in the feeding solution within the FC. Signals were amplified (x200; EX1, Dagan Corporation), digitized with the aid of the A/D converter of the oscilloscope (Tektronix TDS 210) and streamed to a computer for acquisition and visualization (Fig. 1). When the insect inserts its stylets into the feeder and contacts the solution, the electrical circuit closes, and the EMG signals from the suction muscles are recorded. Signals were sampled at 250 Hz.

**Figure 1.**
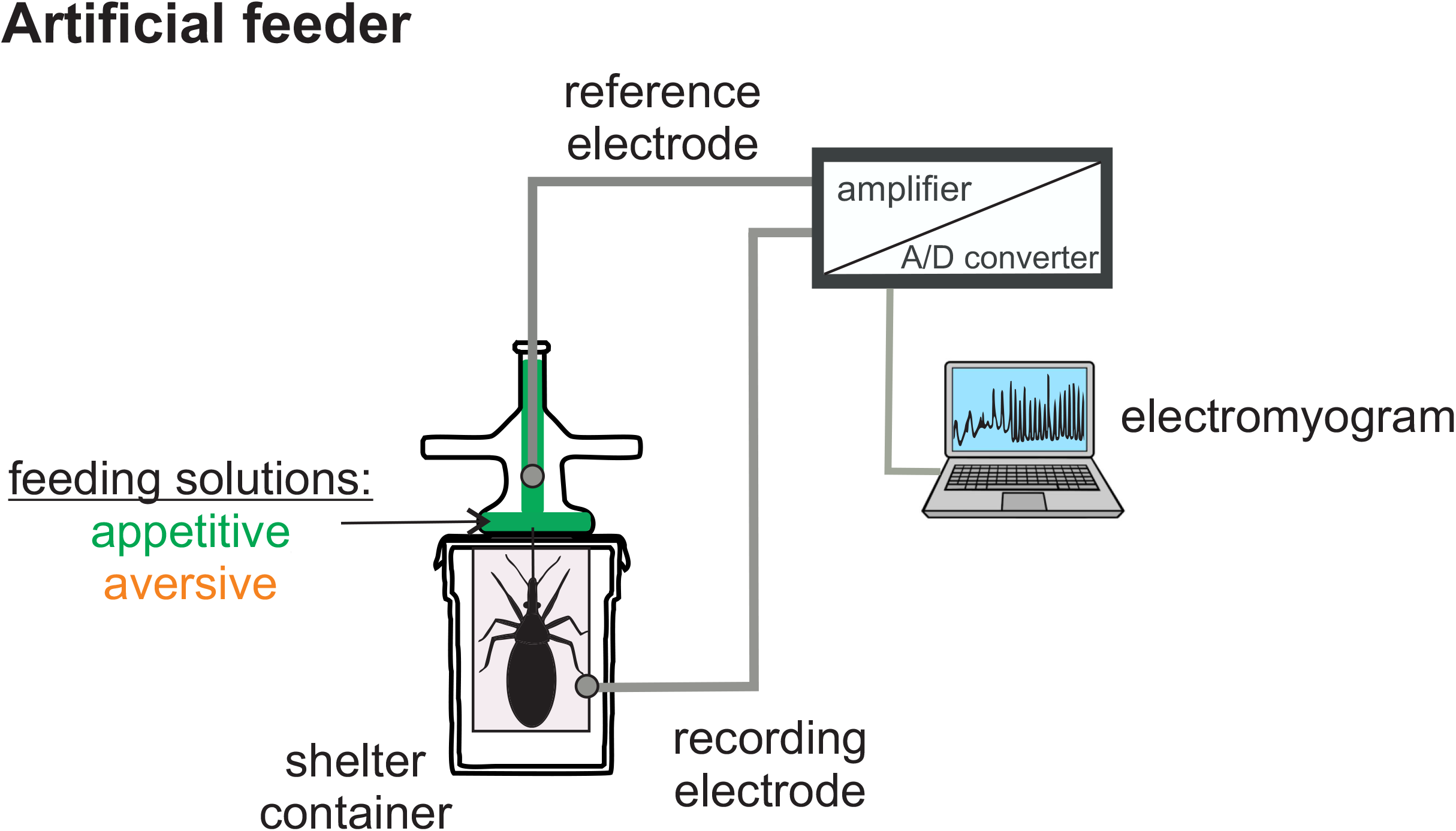
Experimental setup for electromyographic (EMG) recording of feeding behavior. The feeding container (FC) holds the feeding solution and is sealed with a latex membrane simulating host skin. The shelter container (SC), located below the FC, confines the insect. Temperature is maintained at 34 °C using a thermostated bath. A recording electrode is connected to a metal mesh inside the SC, and a reference electrode in the feeding solution within the FC. EMG signals generated during feeding are amplified, digitized and stored on a computer.

### 2.3 Feeding solutions

Appetitive and aversive feeding solutions were used in the experiments to generate a wide range of feeding responses for model training. All solutions were prepared immediately before the experiments and contained 0.15 M NaCl and 1 mM adenosine 5′-triphosphate disodium salt hydrate (ATP) as the base solution, as ATP is a known phagostimulant for blood-feeding insects.

Appetitive stimuli included ATP (0.01, 0.1 and 1 mM; CAS: 56-65-5), BzATP triethylammonium salt (0.1 mM; CAS: 112898-15-4), glucose (10 mM; CAS: CAS 50-99-7), leucine (0.33 mM; CAS: 61-90-5), and phenylalanine (0.33 mM; CAS: 63-91-2). Aversive compounds include saponin (1 mM; CAS: 8047-15-2), suramin hexasodium salt (0.1 mM; CAS: 129-46-4), methyl salicylate (1 mM; CAS: 119-36-8), garlic extract (4 g/ml; CAS: 112898-15-4), caffeine dehydrate (1 mM, 5 mM; CAS: 58-08-2) and quinine hydrochloride dihydrate (1 mM, 5 mM; CAS: 6119-47-7).

These chemically diverse stimuli generated variability in EMG profiles associated with different feeding dynamics, enabling robust training of automated detection algorithms.

### 2.4 MyoRec analysis workflow

EMG recordings were analyzed using MyoRec, a software framework developed to automatically identify ingestion activity and quantify feeding dynamics from electrophysiological recordings. The framework consists of three complementary modules: an Ingestion Event Identifier (IEI), which detects ingestion periods (IPs); a Peak Detector (PD), which identifies individual pumping events within ingestion activity; and a Data Calculator (DC), which extracts behavioral parameters from detected events.

Raw EMG recordings were first preprocessed to improve signal quality and reduce baseline fluctuations. This preprocessing step included baseline correction using the BEADS algorithm, a method for estimating and removing baseline drift, followed by signal normalization to standardize amplitude values across recordings [17].

Preprocessed recordings were then analyzed by the IEI module to automatically identify ingestion periods directly from the EMG traces. Within these periods, the PD module detected individual pumping events. The DC module then quantified behavioral variables, including Contact Period (CP; the interval between stylet insertion and withdrawal), Number of Pumps (NP; total pumping events during ingestion), Duration of Ingestion (DI), Pumping Frequency (PF = NP/DI), Sampling Duration (SD), and Peak Amplitude (A). Pumping frequency helps distinguish periods of active ingestion from pauses during feeder contact.

The framework supports both supervised inspection and fully automated batch processing of EMG recordings.

### 2.5 Automated detection of ingestion activity

Different signal representations were evaluated for automated detection of ingestion activity, including raw EMG signals, Fast Fourier Transform (FFT)-transformed signals, and combined raw+FFT representations.

Multiple machine-learning approaches were evaluated for ingestion event detection, including logistic regression (LR), support vector machines (SVM), gradient boosting (GB), random forest (RF), recurrent neural network (RNN), long short-term memory network (LSTM), convolutional neural networks (CNNt1 and CNNt2), and ensemble methods. These models were trained and evaluated for their ability to automatically identify ingestion periods from EMG recordings, with performance assessed using cross-validation and standard classification metrics. Additional details regarding model architecture, training procedures, and segmentation refinement are provided in the Supplementary Methods.

### 2.6 Model training and validation

A total of 60 EMG recordings were used to create the training dataset, producing 27,376 time windows. Datasets were balanced using undersampling to prevent classification bias. The data were split into 80% for training and validation and 20% for testing. Model performance was evaluated using ten-fold cross-validation, with accuracy as the primary classification metric (see Supplementary Material).

### 2.7 Algorithm performance evaluation

The performance of the Data Calculator module was evaluated by comparing the automated extraction of feeding parameters against reference measurements obtained using the semi-supervised Peak Detector module. A total of 114 EMG recordings were analyzed.

For each recording, parameter extraction accuracy was evaluated by comparing pumping frequency and peak amplitude values automatically computed by the Data Calculator (DC) to reference measurements obtained through supervised analysis using the Peak Detector (PD) workflow. Error rates were quantified using the weighted mean absolute percentage error (WMAPE), which estimates the average percentage deviation between automated and reference measurements while accounting for differences in signal magnitude across recordings. Detailed validation procedures are provided in the Supplementary Methods.

### 2.8 Statistical analysis

Behavioral variables obtained under appetitive and aversive conditions were compared using the non-parametric Mann–Whitney U test. Statistical analyses were performed for contact period duration (CP), sampling phase duration (SP), ingestion duration (DI), pumping frequency (PF), and number of pumping events (NP). Differences were considered statistically significant at α = 0.05.

## 3. Results

### 3.1 Electromyographic signatures of feeding behavior

Electromyographic (EMG) recordings revealed distinct patterns associated with different phases of feeding behavior in *Rhodnius prolixus* (Fig. 2). Feeding behavior during contact with the artificial feeder is organized into a Contact Period (CP), defined as the interval between stylet insertion and withdrawal. Within each CP, insects may perform brief intake events referred to as the Sampling Phase (SP), characterized by irregular, low-amplitude pumping activity associated with food evaluation. If feeding proceeds, insects transition to an Ingestion Phase (IP), characterized by sustained, regular, high-amplitude pumping activity. Multiple CPs, SPs, and IPs can occur within a single recording and are identifiable in EMG traces by distinct voltage patterns.

**Figure 2.**
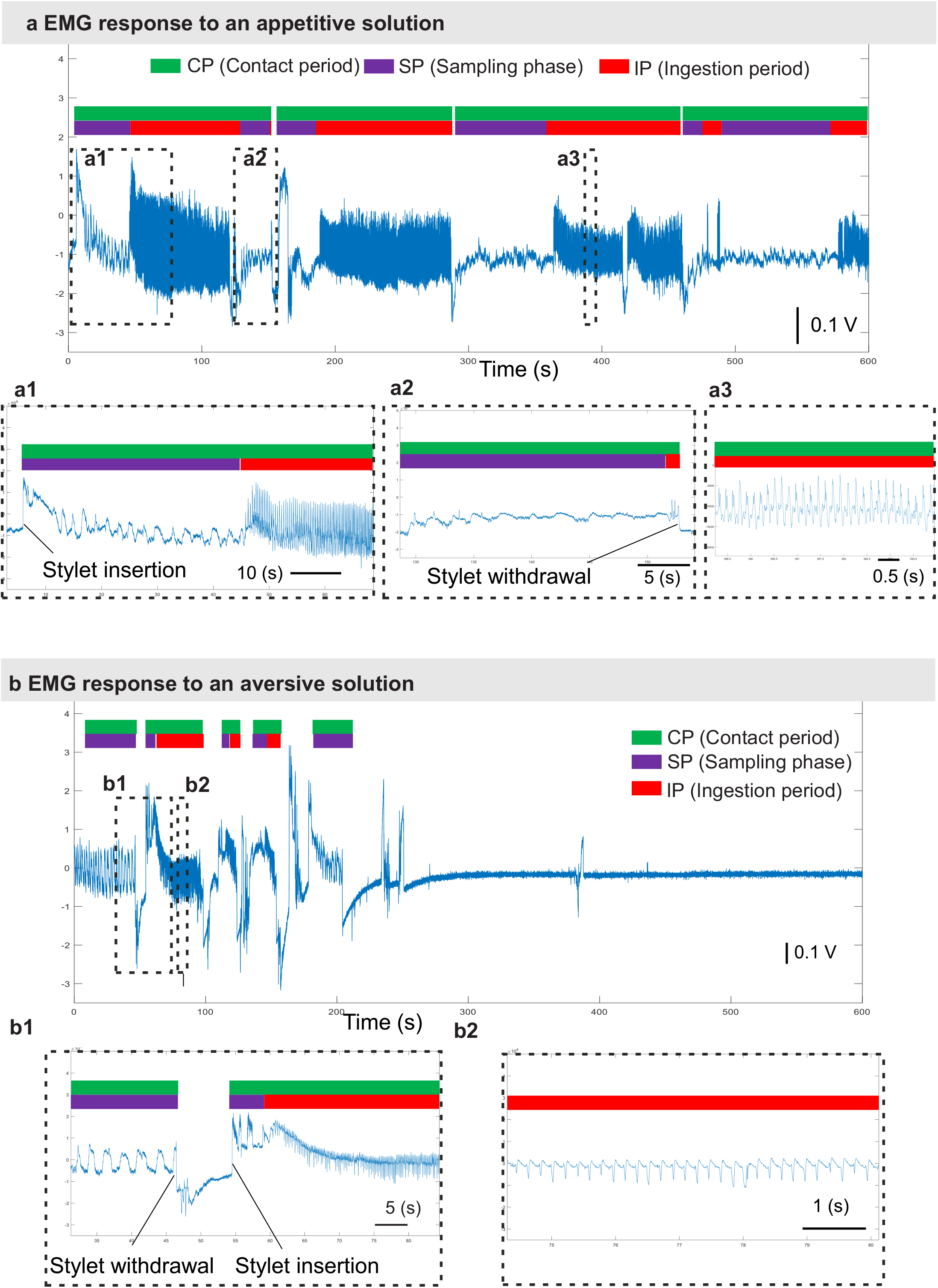
Electromyographic (EMG) patterns associated with feeding behavior. Representative EMGs recordings obtained from insects exposed to appetitive and aversive solutions. Colored bars indicate Contact Period (CP, green), Sampling Phase (SP, purple), and Ingestion Phase (IP, red). **a -** EMG trace from an insect fed on an appetitive ATP-NaCl solution. **a1 -** Expanded view showing stylet insertion and low-amplitude voltage deflections characteristic of the sampling phase (SP), followed by the onset of ingestion (IP). **a2 -** Example of a long sampling phase, followed by a short ingestion phase that ends with the stylet withdrawal. **a3 -** Expanded view showing regular, high-amplitude voltage peaks characteristic of sustained ingestion IP. **b -** EMG trace from an insect fed on an aversive caffeine solution. **b1 -** Expanded view showing stylet withdrawal after a sampling phase (SP), followed by an insertion, a new sampling phase (SP), and an ingestion phase (IP). **b2 -** Example of an ingestion phase (IP).

The structure and duration of these phases varied depending on the gustatory properties of the feeding solution (Fig. 2). Appetitive conditions were typically associated with prolonged ingestion phases and sustained pumping activity (Fig. 2a), whereas aversive conditions were associated with shorter contact periods, fewer pumping events, and rapid termination of ingestion (Fig. 2b). These differences highlight the dynamic organization of feeding behavior and establish a framework for its quantitative analysis.

### 3.2 Automated framework for quantifying feeding dynamics

Although the different phases of feeding behavior can be identified by visual inspection of EMG recordings, systematic quantification across multiple recordings remains labor-intensive and subject to variability. In particular, detecting ingestion phases and extracting pumping dynamics requires precise identification of signal features, which may vary across individuals and experimental conditions.

To enable quantitative analysis of feeding behavior from EMG recordings, we implemented MyoRec, an automated analysis framework that identifies ingestion activity and extracts behavioral parameters directly from EMG signals (Fig. 3a). This framework consists of three components: the Ingestion Event Identifier (IEI) for detecting ingestion phases, the Peak Detector (PD) for identifying individual pumping events, and the Data Calculator (DC) for fully automated extraction of behavioral parameters. These modules allow consistent and reproducible quantification of parameters such as contact period duration, sampling phase duration, ingestion duration, number of pumping events, peak amplitude, and pumping frequency across recordings. The MyoRec workflow includes signal preprocessing (baseline correction and normalization), automated ingestion detection, pumping event identification, and parameter extraction (Fig. 3a). The framework supports both supervised inspection and fully automated batch processing, enabling efficient analysis of large datasets with minimal user intervention.

**Figure 3.**
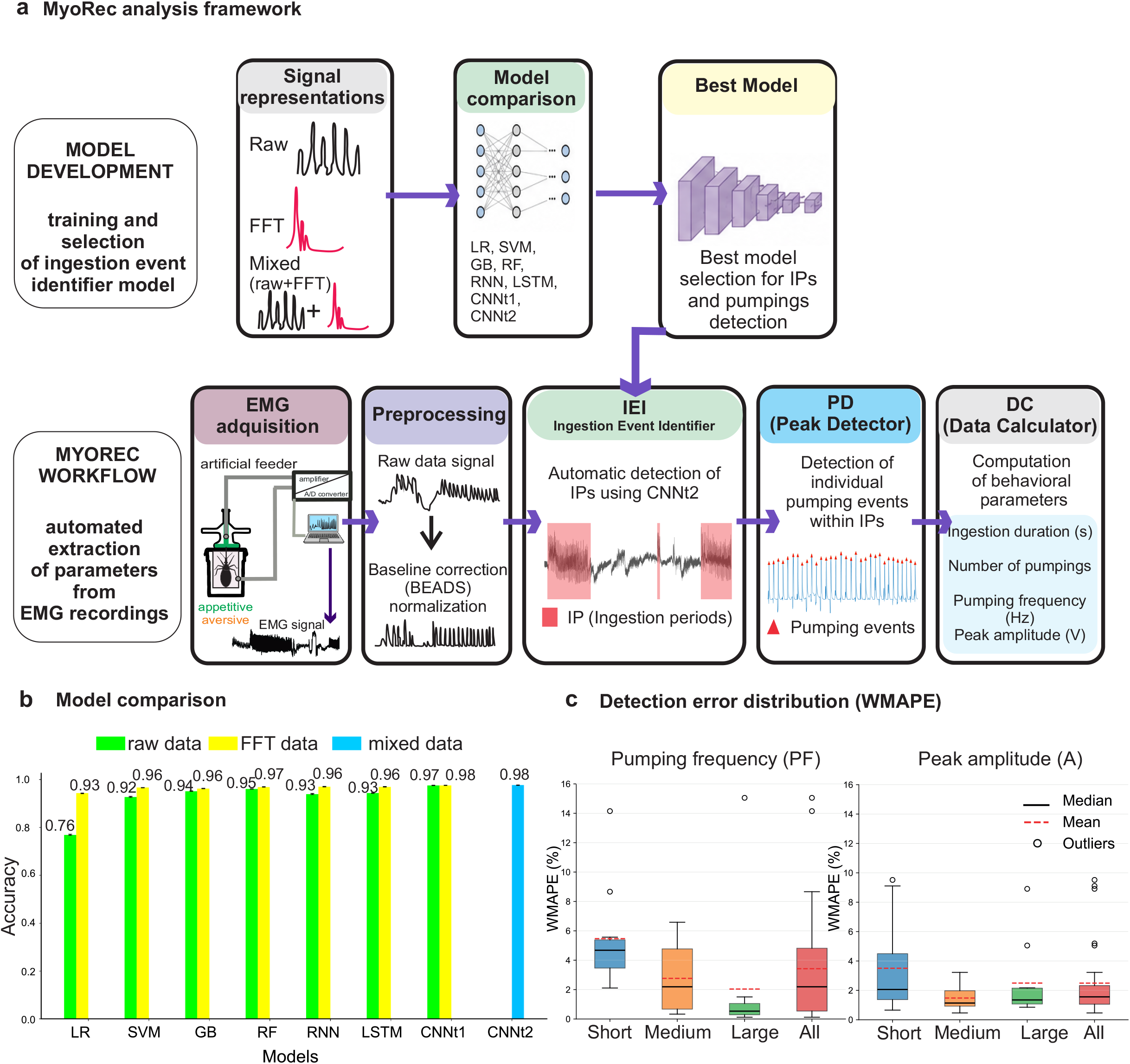
MyoRec framework for automated analysis of EMG recordings. **a -** Overview of the MyoRec framework. The upper panel illustrates the model development stage, including evaluation of different signal representations (raw, FFT, and mixed raw+FFT data) and comparison of multiple machine learning architectures for ingestion event identification. The lower panel shows the operational workflow used for automated analysis of EMG recordings. Raw EMG signals obtained during insect feeding are preprocessed through baseline correction and normalization, followed by automatic identification of ingestion periods (IPs) using the Ingestion Event Identifier (IEI). Individual pumping events within IPs are subsequently detected by the Peak Detector (PD), and behavioral parameters are quantified using the Data Calculator (DC). **b -** Comparison of classification accuracy obtained with different machine learning models and signal representations. Models evaluated included Logistic Regression (LR), Support Vector Machine (SVM), Gradient Boosting (GB), Random Forest (RF), Recurrent Neural Network (RNN), Long Short-Term Memory (LSTM), and convolutional neural networks (CNNt1 and CNNt2). The inclusion of FFT information improved classification performance across models, with convolutional neural networks achieving the highest overall accuracy. CNNt2, which integrates raw and FFT representations, achieved the best performance among the evaluated approaches. **c -** Distribution of weighted mean absolute percentage error (WMAPE) for automated extraction of pumping frequency (PF) and peak amplitude (A) across recordings of different durations. Error values remained low overall and decreased in recordings of intermediate and long duration, indicating strong agreement between automated and supervised measurements.

These capabilities provide a practical solution for scaling the quantitative analysis of feeding dynamics and enable systematic comparisons across experimental conditions.

### 3.3 Identification of ingestion activity from EMG signals

Reliable detection of ingestion activity is essential for quantifying feeding behavior from EMG recordings. We therefore evaluated whether ingestion periods (IPs) could be automatically identified using computational approaches.

EMG signals were analyzed using both raw time-domain data and frequency-domain representations obtained through Fast Fourier Transformation (FFT) (Fig. 3a,b). Across all models, classification performance improved when FFT information was included, indicating that ingestion activity contains rhythmic and frequency-dependent patterns that can be effectively captured in the spectral domain.

Multiple machine learning architectures were evaluated, including Logistic Regression (LR), Support Vector Machine (SVM), Gradient Boosting (GB), Random Forest (RF), Recurrent Neural Network (RNN), Long Short-Term Memory (LSTM), and convolutional neural networks (CNNs). Among the tested approaches, CNN-based models showed the highest overall performance. In particular, CNNt2, which integrates both raw and FFT representations, achieved the highest classification accuracy (97.7% ± 0.0009%), outperforming alternative models (Fig. 3b, Supplementary Fig. S1, Supplementary repository Files: https://github.com/hecsalms/Model-Comparisons.git).

Based on its accuracy and computational efficiency, CNNt2 was selected as the final model for the IEI module (Fig. 3a,b). A refinement algorithm was incorporated to improve segmentation accuracy by correcting boundary errors and reducing false detections in recordings with irregular activity patterns (Supplementary Fig. S2). Although CNNt1 achieved comparable accuracy, its training time was substantially longer (1.78 ± 0.01 min per run) compared to CNNt2 (0.30 ± 0.01 min; Supplementary Fig. S3), due to the requirement for smaller batch sizes.

These results demonstrate that ingestion activity can be robustly identified from EMG signals, providing a reliable basis for automated quantification of feeding behavior.

### 3.4 Extraction and accuracy of behavioral variables from EMG recordings

Having established reliable identification of ingestion periods (IPs), we next evaluated whether quantitative behavioral parameters could be accurately extracted from EMG recordings. Using the automated framework, individual pumping events detected within ingestion periods (Fig. 3a) were used to compute behavioral variables such as pumping frequency (PF) and peak amplitude (A). The high temporal resolution of the framework enables detection of hundreds to thousands of pumping events within seconds to minutes, facilitating efficient analysis of large datasets.

To assess parameter extraction accuracy, PF and A values automatically computed by the Data Calculator (DC) were compared to reference measurements obtained through supervised validation using the Peak Detector (PD) workflow. Error was quantified using the weighted mean absolute percentage error (WMAPE) across recordings of varying durations (Fig. 3c).

Overall, error values were low for both parameters. Short EMG recordings exhibited higher variability (PF: 5.47% ± 3.36%; A: 3.50% ± 3.13%), likely due to fewer detectable events and reduced signal stability. Error variability decreased with longer recording durations, reaching 2-3% for both PF and A in intermediate and long recordings.

Across all recordings, the overall error remained low (PF: 3.42% ± 3.72%; A: 2.49% ± 2.48%), indicating strong agreement with supervised measurements (Fig. 3c).

These results demonstrate that behavioral parameters can be rapidly, accurately, and reproducibly extracted from EMG recordings using automated analysis, supporting large-scale quantification of feeding dynamics.

### 3.4 Quantifying feeding behavior under different gustatory conditions

Finally, we applied MyoRec to quantify the feeding behavior of *R. prolixus* exposed to solutions with either appetitive or aversive gustatory properties (Fig. 4).

**Figure 4.**
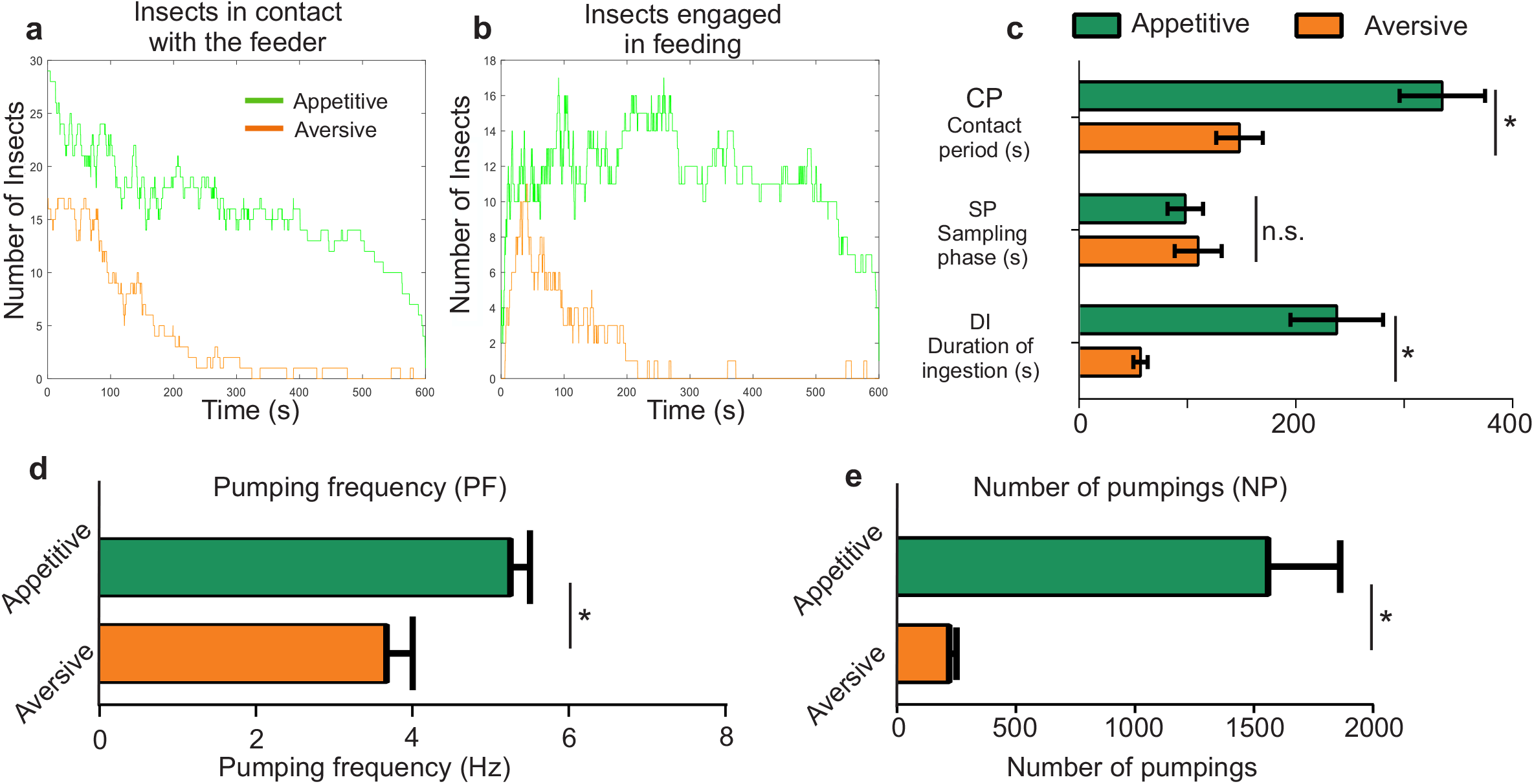
Quantitative analysis of feeding behavior under appetitive and aversive gustatory conditions. **a -** Number of insects maintaining contact with the feeder over time under appetitive (green) and aversive (orange) conditions. Appetitive solutions promoted prolonged contact with the feeder, whereas aversive solutions induced rapid disengagement. **b -** Number of insects actively engaged in ingestion over time. Sustained ingestion activity was observed under appetitive conditions (green), while ingestion rapidly declined under aversive conditions (orange). **c -** Quantification of feeding phases. Contact period duration (CP) and ingestion duration (DI) were significantly longer in insects exposed to appetitive solutions, whereas sampling phase duration (SP) did not differ significantly between treatments. **d -** Pumping frequency (PF) during ingestion. Appetitive solutions elicited higher pumping frequencies than aversive solutions. **e -** Total number of pumping events (NP). Insects exposed to appetitive solutions performed substantially more pumping events than insects exposed to aversive solutions. Error bars indicate SEM. Asterisks indicate statistically significant differences (*p < 0.05); n.s., non-significant.

Clear differences emerged between treatments (Fig. 4). Insects exposed to appetitive solutions maintained prolonged contact with the feeder and sustained ingestion activity over time, whereas insects exposed to aversive solutions rapidly interrupted feeding and disengaged shortly after stylet insertion (Fig. 4a,b). These results indicate that gustatory cues influence both the persistence of feeder contact and the maintenance of ingestion behavior.

Quantitative analysis confirmed these differences. Contact period duration (CP) and ingestion duration (DI) were significantly longer in insects exposed to appetitive solutions than in those exposed to aversive solutions (p < 0.05 for both; Fig. 4c). In contrast, sampling phase duration (SP) did not differ significantly between treatments (n.s.), suggesting that insects initially evaluate both food sources similarly before divergent feeding decisions emerge during ingestion.

Appetitive solutions also elicited significantly higher pumping frequency (PF) and a greater number of pumping events (NP) compared to aversive solutions (p < 0.05 for both; Fig. 4d,e). Together, these results indicate that gustatory valence modulates not only feeding persistence, but also the dynamics of pumping activity during ingestion.

Overall, these findings show that the sensory properties of the feeding solution are associated with differences in feeding decisions and ingestion maintenance, and demonstrate that MyoRec can quantitatively resolve these behavioral processes with high temporal precision.

## 4. Discussion

Feeding in hematophagous insects involves a sequence of behavioral decisions, with insects evaluating the chemical composition of ingested fluids before committing to sustained ingestion. Early electromyographic (EMG) studies in blood-feeding arthropods demonstrated that feeding behavior consists of distinct and quantifiable motor patterns [9,18,19]. Since then, EMG recordings have provided a direct physiological readout of feeding dynamics and ingestion activity, but analysis has traditionally relied on manual annotation and supervised inspection, limiting the scalability, speed, and reproducibility.

In this study, we developed an automated framework that enables reliable identification of ingestion activity and the extraction of quantitative behavioral parameters from EMG recordings in the triatomine *R. prolixus*. By combining signal analysis with machine learning approaches, this framework enables the characterization of feeding dynamics at high temporal resolution while substantially reducing data processing time.

Automated analysis of feeding-related electrical signals has been extensively developed in the context of electropenetrography (EPG), particularly for phytophagous insects such as aphids [20–26], and more recently in mosquitoes [27]. These approaches have enabled the classification of feeding waveforms and the extraction of behavioral parameters from insect-food interactions. In contrast, equivalent tools for EMG recordings in hematophagous insects remain limited. Unlike EPG, which records electrical changes generated during insect– substrate interactions, EMG captures ingestion-related muscular activity directly. The framework presented here adapts the concept of automated signal analysis to a different biological system, providing a tool adapted to the characterization of feeding dynamics in blood-feeding insects.

Our results show that ingestion periods can be reliably identified from EMG signals using automated classification methods. The high detection accuracy indicates that ingestion activity generates consistent temporal and spectral patterns that can be computationally recognized. Notably, detection performance improved with the inclusion of frequency-domain information, reflecting the rhythmic nature of pumping activity during sustained ingestion.

Beyond event detection, automated extraction of behavioral parameters showed strong agreement with supervised analysis across recordings of different durations (Fig. 3c). Although error values were higher in short recordings, likely due to reduced signal stability and lower numbers of detectable pumping events, overall accuracy remained within a range that supports reliable quantitative analysis. These results indicate that automated approaches can reproduce manual or supervised measurements while enabling large-scale analysis that would otherwise be impractical using manual annotation alone.

When applied to insects exposed to stimuli with opposite gustatory valence, marked differences in feeding dynamics were observed. Appetitive solutions promoted sustained ingestion and prolonged contact with the feeder, while aversive conditions led to rapid interruption of feeding and reduced ingestion activity. Notably, sampling phase duration did not differ significantly between treatments, whereas ingestion duration and pumping activity were strongly affected. This suggests that insects initially evaluate both food sources similarly, but diverge during the maintenance of ingestion as sensory information accumulates. These findings are consistent with previous studies showing that gustatory cues regulate feeding decisions in hematophagous insects and demonstrate that such effects can be quantified with high temporal resolution [10,12,13,28]. In addition to altering ingestion persistence, gustatory valence modulated the dynamics of pumping activity, affecting both frequency and the total number of pumping events. These results highlight the dynamic nature of feeding behavior, where both initiation and maintenance of ingestion are continuously modulated by gustatory input, and underscore the value of quantitative approaches for investigating feeding decisions at fine temporal scales.

The framework presented here was developed using a diverse set of chemical stimuli to generate variability in feeding responses, enabling robust training and evaluation of detection algorithms. While this study establishes a general approach to quantifying feeding dynamics, future work could systematically compare the effects of specific compounds to identify gustatory mechanisms underlying feeding modulation.

More broadly, automated analysis of EMG recordings may be applicable to other blood-feeding arthropods and could facilitate comparative studies of feeding physiology across species. By providing quantitative resolution of ingestion dynamics at high temporal resolution, this framework provides new opportunities to investigate how gustatory information shapes the transition from food evaluation to sustained feeding behavior in blood-feeding insects.

## 5. Conclusions

In this study, we developed MyoRec, an automated framework for the detection and quantitative analysis of feeding behavior from EMG recordings in *R. prolixus*. By enabling reliable identification of ingestion activity and accurate extraction of behavioral parameters, this approach substantially reduces analysis time while maintaining consistency with supervised methods.

Application of this framework revealed that gustatory cues dynamically shape feeding behavior, primarily by modulating the maintenance and persistence of ingestion rather than the initial evaluation of food sources. These findings demonstrate that automated analysis of EMG signals provides a practical and scalable tool for investigating feeding decisions in hematophagous insects and offers new opportunities to quantitatively link sensory input with behavioral output at high temporal resolution.

## Acknowledgements

The authors thank the financial support from Agencia Nacional de Promoción de la Investigación, el Desarrollo Tecnológico y la Innovación (Agencia I+D+i) (PICT-2019-02057 to RB); and the Consejo Nacional de Investigaciones Científicas y Técnicas (CONICET) fellowship for Latin American students (granted to HSM); CONICET (PIP 11220220100596CO to RBB), INEE-CNRS (REPEL IRP project).

## Author Contributions

Conceived and designed the project: R.B.B, P.E.S. Data acquisition: I.O.I, R.B.B. Analyzed the data: H.S.M., I.O.I., M.G.A., A.M. Wrote the manuscript: H.S.M., P.E.S, R.B.B. Software Development: H.S.M., M.G.A., A.M.

## Data Availability

MyoRec is freely available at [https://github.com/hecsalms/MyoRec_Dist_Version], together with documentation and example datasets to facilitate reproducibility.

