## Supplementary material for "Automated analysis of feeding dynamics from electromyographic recordings in a blood-sucking insect": Suppl Material

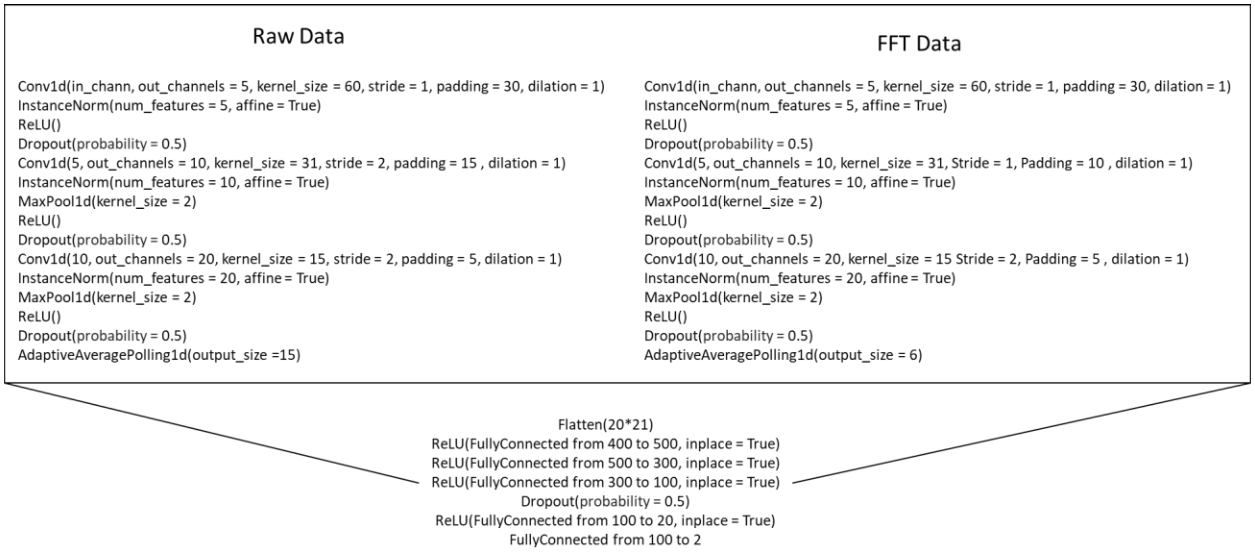


**Supplementary Fig. S1. Architecture of the Type II convolutional neural network (CNNt2).**

The model receives EMG signal segments as input and classifies them into two categories: ingestion-related activity (with peaks) and non-ingestion activity (without peaks). This architecture integrates temporal and frequency-domain features and was selected for the ingestion event identifier (IEI) based on its superior performance.

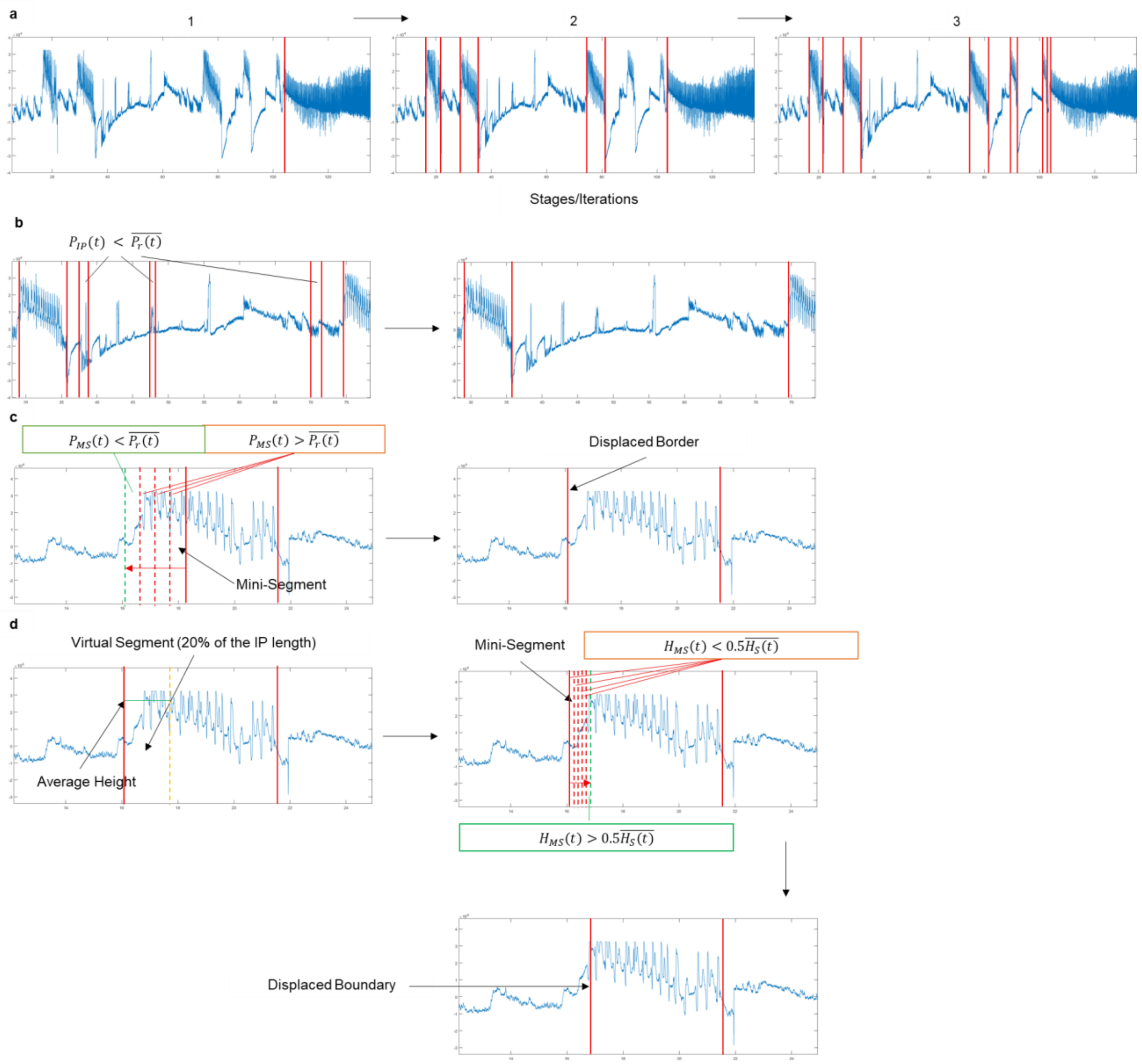


**Supplementary Fig. S2. Refinement procedure for ingestion phase detection. .**

(a) Iterative classification using CNNt2 with multiple window sizes to detect ingestion phases (IPs) of different durations.

(b) Removal of false positives: IPs shorter than 4 s and with signal power below the average are discarded.

(c) Boundary adjustment based on signal power in local mini-segments (MS): boundaries are extended when MS power exceeds the average power of the ingestion phase.

(d) Boundary correction using peak detection: peaks amplitudes within a virtual segment (VS; 20% of IP length) are compared with local maxima, and boundaries are adjusted when peak amplitude criteria are not met.

In panels (c–d), opposite conditions must be satisfied for boundary contraction.

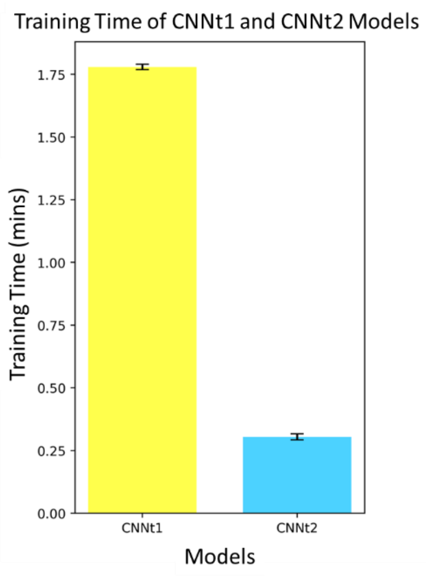


**Supplementary Fig. S3. Training time of CNN models.**

Training time (in minutes) for CNNt1 and CNNt2 models. CNNt2 achieves comparable accuracy with substantially reduced training time.

**Table S1 Optimized hyperparameters for machine learning models trained on EMG data.**

The table shows the main hyperparameters selected for each model after optimization using automated and manual tuning procedures. Models were trained using either raw EMG signals (Raw) or frequency-domain features obtained by Fast Fourier Transform (FFT). Only the most relevant hyperparameters are reported.

| **Model** | **Hyperparameter** | **Library** | **Raw** | **FFT** |
|---|---|---|---|---|
| Logistic Regression (LR) | C | cuML | 50.75 | 231.59 |
| | max_iter | | 239 | 1310 |
| Support Vector Machines (SVM) | C | ThunderSVM | 9.19 | 8.67 |
| | gamma | | 0.02 | 0.15 |
| Random Forest (RF) | max_depth | cuML | 30 | 21 |
| | n_estimators | | 127 | 108 |
| Gradient Boosting (GB) | learning_rate | CatBoost | 0.20 | 0.14 |
| | max_depth | | 7 | 5 |
| | n_estimators | | 250 | 260 |
| Recurrent Neural Network (RNN) | optimizer__lr | Pytorch-Skorch | 0.01 | 0.01 |
| | batch_size | | 200 | 200 |
| | max_epochs | | 80 | 80 |
| Long Short-Term Neural Network (LSTM) | optimizer__lr | Pytorch-Skorch | 0.001 | 0.01 |
| | batch_size | | 200 | 200 |

| | | | | |
|---|---|---|---|---|
| | max_epochs | | 80 | 70 |
