## Supplementary material for "Automated analysis of feeding dynamics from electromyographic recordings in a blood-sucking insect": Suppl Methods

### Supplementary Methods

#### Signal preprocessing

Prior to segmentation and model training, electromyographic (EMG) recordings were preprocessed to improve signal quality and facilitate automated pattern recognition. Baseline correction and denoising were performed using the Base Line Estimation and Denoising With Sparsity (BEADS) algorithm (Ning et al., 2014). This method removes low-frequency trends and irregular baseline fluctuations while preserving sharp transient features such as voltage peaks associated with cibarial pumping activity. By correcting baseline drift and reducing noise, the BEADS algorithm improves the signal-to-noise ratio and enhances the detection of waveform patterns used for model training.

#### Dataset generation and feature extraction

Raw EMG recordings were segmented into sequential one-second windows. Each window contained 250 samples, corresponding to the acquisition sampling rate of 250 Hz. Each segmented window was treated as an independent instance for machine-learning analysis.

Two feature representations were generated from each EMG window. The first consisted of the original voltage values from the raw signal. The second representation was obtained by applying the Fast Fourier Transform (FFT) to each window, generating a set of frequency-domain features that describe the spectral composition of the signal. The FFT transformation produced 126 frequency components per instance.

These two representations allowed models to learn patterns in both the time domain and the frequency domain.

#### Instance labeling and dataset balancing

Training instances were labeled according to manually identified ingestion phases in the EMG recordings. Manual annotation of ingestion phases was performed using visualization tools implemented in the MyoRec software.

Each segmented window was assigned a class label depending on its overlap with annotated ingestion phases. Windows whose duration overlapped by at least 50% with a manually defined ingestion phase were labeled as "With Peaks", while all other windows were labeled as "Without Peaks".

To prevent classification bias caused by class imbalance, datasets were balanced using an undersampling approach. Instances belonging to the majority class ("Without Peaks") were randomly reduced until both classes contained equal numbers of samples.

#### Machine learning models evaluated

Several machine learning models were evaluated for ingestion-event detection. The tested models included logistic regression, decision trees, support vector machines (SVM), and ensemble methods such as random forest and gradient boosting.

Deep learning architectures were also evaluated, including convolutional neural networks (CNN), recurrent neural networks (RNN), and long short-term memory networks (LSTM). Both individual models and stacked model architectures were tested to determine which approach provided the highest predictive performance.

#### Convolutional neural network architectures

Two convolutional neural network architectures were implemented. The first architecture (CNN Type I, hereafter CNNt1) consisted of a relatively simple configuration with a small number of convolutional layers designed to capture local signal patterns while maintaining computational efficiency.

The second architecture (CNN Type II, hereafter CNNt2) was designed with a deeper structure including additional convolutional layers. This architecture received two input streams simultaneously: raw EMG signal segments and FFT-derived frequency-domain features. By combining time-domain and frequency-domain information, the model learned complementary signal features associated with ingestion activity (see Supplementary Fig. S1).

Stacked models combining CNNt2 predictions with additional classifiers, including random forest, gradient boosting, SVM, and LSTM networks, were also evaluated.

**Hyperparameter optimization**

Hyperparameters for all tested models were optimized using a combination of automated and manual approaches (see Supplementary Table S1). Automated optimization methods included Tree-structured Parzen Estimators (TPE), grid search, and randomized search. These procedures explored different combinations of parameters in order to maximize classification performance.

After automated optimization, additional manual tuning was performed to refine model architecture and improve predictive accuracy.

**Refinement algorithm for ingestion phase detection**

To improve the accuracy of ingestion phase detection and reduce classification errors, a refinement algorithm was applied to the predictions generated by the Ingestion Event Identifier (IEI). For each EMG recording, three input data sizes (segmentation windows) were assumed: large = 750 (Raw), 376 (FFT); medium = 125 (Raw), 63 (FFT); and short = 64 (Raw), 32 (FFT) samples. Thus, the model produced three prediction outputs to identify large, medium and short ingestion phases respectively in the EMGs.

Large inputs were used to detect long ingestion phases (>10 s), medium inputs were used to detect ingestion phases of intermediate duration (3–10 s), and short inputs were used to detect brief ingestion events (<3 s). Final ingestion phase boundaries were determined by overlaying predictions obtained from these input window sizes and retaining non-embedded borders.

An ingestion phase was considered valid when three consecutive windows were classified as containing ingestion activity.

**Signal power filtering**

To eliminate false positive detections, signal power filtering was applied. Signal power was calculated as the average squared amplitude of the signal samples within a given segment, as shown in the following expression:

$$\underline{P\ \ (t)} = \frac{1}{t_2 - t_1 + 1} \sum_{t_1}^{t_2} |x(t)|^2$$

(1)

where $\underline{P\ \ (t)}$ is the threshold power (or average power) of the signal segment (EMG) with limits $t_2$ and $t_1$, while $x(t)$ are the samples it contains.

The average power of ingestion phases longer than four seconds was used as a reference value. Segments whose signal power fell below a user-defined fraction of this reference value were classified as false positives and removed from the final ingestion-phase detection (see Supplementary Fig. S2).

**Border refinement procedure**

Following ingestion-phase detection, phase boundaries were refined using local comparisons of signal power and peak amplitude. Local signal segments surrounding each detected boundary were analyzed iteratively to determine whether boundaries should be extended or reduced.

Signal power within mini-segments surrounding each boundary was compared with threshold values derived from the average signal power of previously identified ingestion phases. Boundary adjustments were performed until the power condition changed or a maximum displacement limit was reached.

An additional and equivalent mechanism used a fraction of the average peak amplitudes within regions corresponding to 20% of the ingestion phase length as thresholds. Average amplitudes were compared to the highest potential values within surrounding mini-segments to adjust boundaries.

**Peak detection algorithm**

Voltage peaks corresponding to pumping events were detected using the Downward Zero-Crossings Smoothed First Derivative (DZCSFD) method (O'Haver, 2025). A signal sample was considered a peak when three conditions were satisfied: the signal amplitude exceeded a predefined threshold, the first derivative of the signal crossed zero, and the second derivative exceeded a predefined slope threshold.

Two peak detection modes were implemented. In the Full Signal Threshold mode, a constant amplitude threshold was applied across the entire ingestion phase. In the Divided Signal Threshold mode, ingestion phases were divided into smaller segments and thresholds were dynamically calculated based on the distribution of peak amplitudes within each segment.

**Peak amplitude correction and filtering**

Because signal smoothing may reduce measured peak amplitudes, a variance-factor correction was implemented. This procedure identifies a local interval surrounding each detected peak and recalculates its amplitude using the original unsmoothed signal.

To avoid interference from neighboring peaks, a reduced interval was applied to peaks with amplitudes smaller than one third of the maximum peak amplitude detected within the ingestion phase.

A horizontal filtering step was also implemented to eliminate small peaks associated with signal noise. Within each defined horizontal interval, only the highest peak was retained.

**Spectral estimation of pumping frequency**

Pumping frequency was estimated using spectral analysis of EMG signals. Power spectral density was calculated using the Welch method applied to overlapping signal segments. The fundamental frequency identified in the frequency spectral domain as the highest peak, was assumed to correspond to the pumping frequency of the cibarial pump.

For short ingestion phases, pumping frequency estimates obtained through spectral analysis were combined with peak-counting estimates in order to improve accuracy.

**Batch parameter estimation**

The Data Calculator module enables automated batch analysis of EMG recordings using fixed parameter configurations. This module calculates behavioral variables for multiple recordings simultaneously.

Three amplitude estimation modes were implemented to allow users to adjust the balance between processing speed and estimation accuracy. Full Amplitude Estimation calculates peak amplitudes across the entire ingestion phase, Partial Amplitude Estimation calculates amplitudes in selected segments of ingestion phases, and No Amplitude Estimation disables amplitude calculations to maximize processing speed.

**Data Calculator: algorithm validation**

The performance of automated parameter extraction was evaluated using a dataset of 114 EMG recordings. Recordings were categorized into three feeding profiles based on the number of pumping events detected: short ingestion profiles (10–100 pump events), medium profiles (100–500 pump events), and large profiles (>500 pump events). To find the most effective hyperparameter configuration for the algorithm (training procedure), 84 recordings were selected and analyzed with the Peaks Detector. A test set with 10 recordings for each profile category was created to compare the performance of the Data Calculator's algorithm to the Peaks Detector one. Recordings with less than 10 pump events were dismissed.

Error rates for pumping frequency and peak amplitude estimates (compared variables), were calculated using the Weighted Mean Absolute Percentage Error (WMAPE) metric, which accounts for the relative contribution of each ingestion phase according to its number of pumping events. It is defined according to the following expression:

$$WMAPE = \frac{1}{N}\sum_{j=1}^{N} \left[\frac{1}{n_j}\sum_{i=1}^{n_j} \left(\frac{|PD_{ij}-DC_{ij}|}{PDij}\right) \times 100 \times W_{ij}\right], j = 1,\ldots,N;\ i = 1,\ldots,n_j$$

(2)

where $WMAPE$ represents the mean weighted percentage error for all records, $PD_{ij}$ and $DC_{ij}$ the PF/A value of the i-th IP of the j-th record calculated with the PD/DC respectively, $n_j$ indicates the number of IPs, $N$ is the number of records, and $W_{ij}$ is the i-th weight of the j-th record (number of peaks inside the i-th IP of the j-th record divided by the total number of peaks in that record). Data Calculator hyperparameters configuration was estimated based on data analyzed using the Peak Detector: WL = 1024 samples, OV = 960 samples, AF = 0.05, ST = 4.6094e-08, SW (Smooth Width) = 3, PG (Peak Group) = 2, STY (Smooth Type) = 4, VF = 2, LVF = 4, and HF = 9 (refer to Supplementary repository Files: https://github.com/hecsalms/Model-Comparisons/DC_Parameters.git).
